# Munc13 supports vesicle fusogenicity after disrupting active zone scaffolds and synaptic vesicle docking

**DOI:** 10.1101/2022.04.01.486686

**Authors:** Chao Tan, Giovanni de Nola, Claire Qiao, Cordelia Imig, Nils Brose, Pascal S. Kaeser

## Abstract

Active zones consist of protein scaffolds that are tightly attached to the presynaptic plasma membrane. They dock and prime synaptic vesicles, couple them to Ca^2+^ entry, and target neurotransmitter release to postsynaptic receptor domains. Simultaneous RIM+ELKS ablation disrupts these scaffolds, abolishes vesicle docking and removes active zone-targeted Munc13, but some vesicles remain releasable. This enduring vesicular fusogenicity may be Munc13-independent or be mediated by non-active zone-anchored Munc13. We tested its Munc13-dependence by ablating Munc13-1 and Munc13-2 on top of RIM+ELKS in cultured hippocampal neurons. The hextuple knockout synapses lacked docked vesicles, but other ultrastructural features were near-normal despite the strong genetic manipulation. Removing Munc13 in addition to RIM+ELKS further impaired action potential-evoked release by decreasing the remaining pool of releasable vesicles. We conclude that Munc13 can support some fusogenicity without RIM and ELKS, and that presynaptic recruitment of Munc13, even without active zone-anchoring, suffices to generate some fusion-competent vesicles.

## Introduction

Neurotransmitter release is mediated by synaptic vesicle fusion at presynaptic active zones, and Munc13 proteins have a central role in this process (Brunger et al., 2018; Dittman, 2019; Wojcik and Brose, 2007). In addition to Munc13, active zone scaffolds contain RIM, ELKS, RIM-BP, Bassoon/Piccolo, and Liprin-α. Together, they form a protein machine that controls the speed and precision of synaptic transmission by docking and priming of synaptic vesicles, by organizing their coupling to sites of Ca^2+^ entry, and by targeting transmitter release towards postsynaptic receptor domains (Biederer et al., 2017; Südhof, 2012). Given the molecular complexity of the active zone, understanding its assembly and functions has remained a key challenge in synaptic neuroscience.

Mouse knockout studies identified a major role of Munc13 in vesicle priming (Augustin et al., 1999; Varoqueaux et al., 2002). This process renders vesicles fusion-competent and adds them to the pool of readily releasable vesicles (RRP) that can rapidly undergo exocytosis upon action potential arrival. The functional RRP can be probed experimentally by applying stimuli that deplete it, for example superfusion of hypotonic sucrose (Kaeser and Regehr, 2017; Rosenmund and Stevens, 1996). When Munc13 is deleted, release-competence of vesicles is abolished or strongly decreased across all tested synapses in multiple model organisms, including mouse, Drosophila melanogaster and Caenorhabditis elegans (Aravamudan et al., 1999; Augustin et al., 1999; Richmond et al., 1999; Varoqueaux et al., 2002). Munc13 ablation also results in an almost complete loss of docked vesicles, as defined by plasma membrane attachment in electron micrographs (Hammarlund et al., 2007; Imig et al., 2014; Siksou et al., 2009). These findings led to the model that vesicle docking and priming are morphological and functional states that correspond to release competence, which is further supported by similar correlations upon ablation of SNARE proteins (Chen et al., 2021; Hammarlund et al., 2007; Imig et al., 2014; Kaeser and Regehr, 2017). Due to their core function in vesicle priming, Munc13 mediates the resupply of fusion-competent vesicles as they are spent during synaptic activity and thereby controls short-term plasticity and recovery from depression (Lipstein et al., 2013, 2021; Rosenmund et al., 2002). Furthermore, Munc13 nano-assemblies may account for secretory hotspots and recruit the SNARE protein syntaxin-1 (Reddy-Alla et al., 2017; Sakamoto et al., 2018).

Several recent studies took the approach to ablate combinations of active zone protein families to analyze active zone assembly and function by eliminating redundant components (Acuna et al., 2016; Brockmann et al., 2020; Kushibiki et al., 2019; Oh et al., 2021; Tan et al., 2022; Wang et al., 2016; Zarebidaki et al., 2020). We simultaneously deleted RIM1, RIM2, ELKS1 and ELKS2, which resulted in a loss of RIM, ELKS, and Munc13, and in a strong decrease of Bassoon, Piccolo, and RIM-BP levels at active zones (Tan et al., 2022; Wang et al., 2016; Wong et al., 2018). This disruption of active zone assembly led to a near-complete loss of synaptic vesicle docking as studied by electron microscopy and to an ∼85% decrease in single action potential-evoked exocytosis as assessed electrophysiologically. However, some transmitter release persisted: up to ∼35% of release evoked by hypertonic sucrose and ∼50% of spontaneous release events remained, increasing extracellular Ca^2+^ to enhance vesicular release probability p strongly enhanced single action potential-evoked release, and stimulus trains released vesicles quite efficiently (Wang et al., 2016). Hence, some releasable vesicles persisted despite the loss of most docked vesicles. These findings led to alternative mechanistic models in which docking and priming are independent processes mediated by distinct molecular functions of Munc13 (Kaeser and Regehr, 2017). Further support for this model came from experiments with artificial re-targeting of Munc13 to synaptic vesicles rather than the active zone, which increases vesicle fusogenicity but not docking in mutants that lack most RIM and ELKS sequences and docked vesicles (Tan et al., 2022). Together, these findings indicate that non-docked vesicles can contribute to the functional RRP after removal of RIM and ELKS. What remained enigmatic, particularly in view of the near-complete loss of Munc13 from the target membrane after RIM+ELKS ablation, was whether undocked vesicles engaged Munc13 for vesicle priming or used an alternative priming pathway.

In the present study, we tested directly whether Munc13 is sufficient to prime vesicles after the strong active zone disruption upon RIM+ELKS knockout. We generated mice for ablating Munc13-1 and Munc13-2 in addition to RIM and ELKS (that is RIM1, RIM2, ELKS1 and ELKS2) and compared phenotypes of hextuple knockout neurons generated from these mice with those from conditional RIM+ELKS knockouts only. We find that synapses form despite this strong genetic manipulation and that overall, their ultrastructure is largely normal except for a lack of docked vesicles. However, Munc13 ablation on top of RIM and ELKS knockout further impaired single action potential-evoked release and decreased the RRP at both excitatory and inhibitory synapses. Paired pulse ratios, used to monitor p, were not further affected by the additional removal of Munc13. Our data establish that Munc13 can functionally prime some vesicles in the absence of RIM and ELKS, indicate that Munc13 away from active zones is sufficient to enhance vesicle fusogenicity, and support a growing body of data showing that synapse formation is overall remarkably resilient to severe perturbations of synaptic protein content and of synaptic activity. We propose that Munc13 recruitment to presynapses is rate-limiting to generate fusion competence of synaptic vesicles.

## Results

### Some synaptic Munc13 remains after RIM+ELKS knockout

With the overall goal to determine whether Munc13 mediates addition of vesicles to the functional RRP after RIM+ELKS knockout, we first assessed key effects on synaptic transmission and Munc13 active zone levels in cultured neurons after ablation of RIM+ELKS (Figs. 1A-1J). We cultured primary hippocampal neurons of mice with floxed alleles for RIM1, RIM2, ELKS1 and ELKS2 (Fig. 1A) as described before (Tan et al., 2022; Wang et al., 2016). At 5 days in vitro (DIV), the neurons were transduced with Cre-expressing lentiviruses or control lentiviruses (that express an inactive mutant of cre) to generate cKO^R+E^ neurons or control^R+E^ neurons, respectively. We previously established that this induces strong defects in active zone assembly and neurotransmitter release, but the neurons form synapses at normal numbers, and postsynaptic receptor assemblies and functions are preserved (Tan et al., 2022; Wang et al., 2016). We first confirmed that excitatory and inhibitory synaptic transmission are strongly impaired in cKO^R+E^ neurons (Figs. 1B-1F). Synaptic responses were induced by focal electrical stimulation, and whole cell recordings served to monitor excitatory and inhibitory transmission via glutamate and GABA_A_ receptors, respectively. Action potential-evoked excitatory transmission was monitored via NMDA receptors to prevent the strong reverberant activity that is induced by stimulation of these neuronal networks when AMPARs are not blocked.

**Figure 1.**
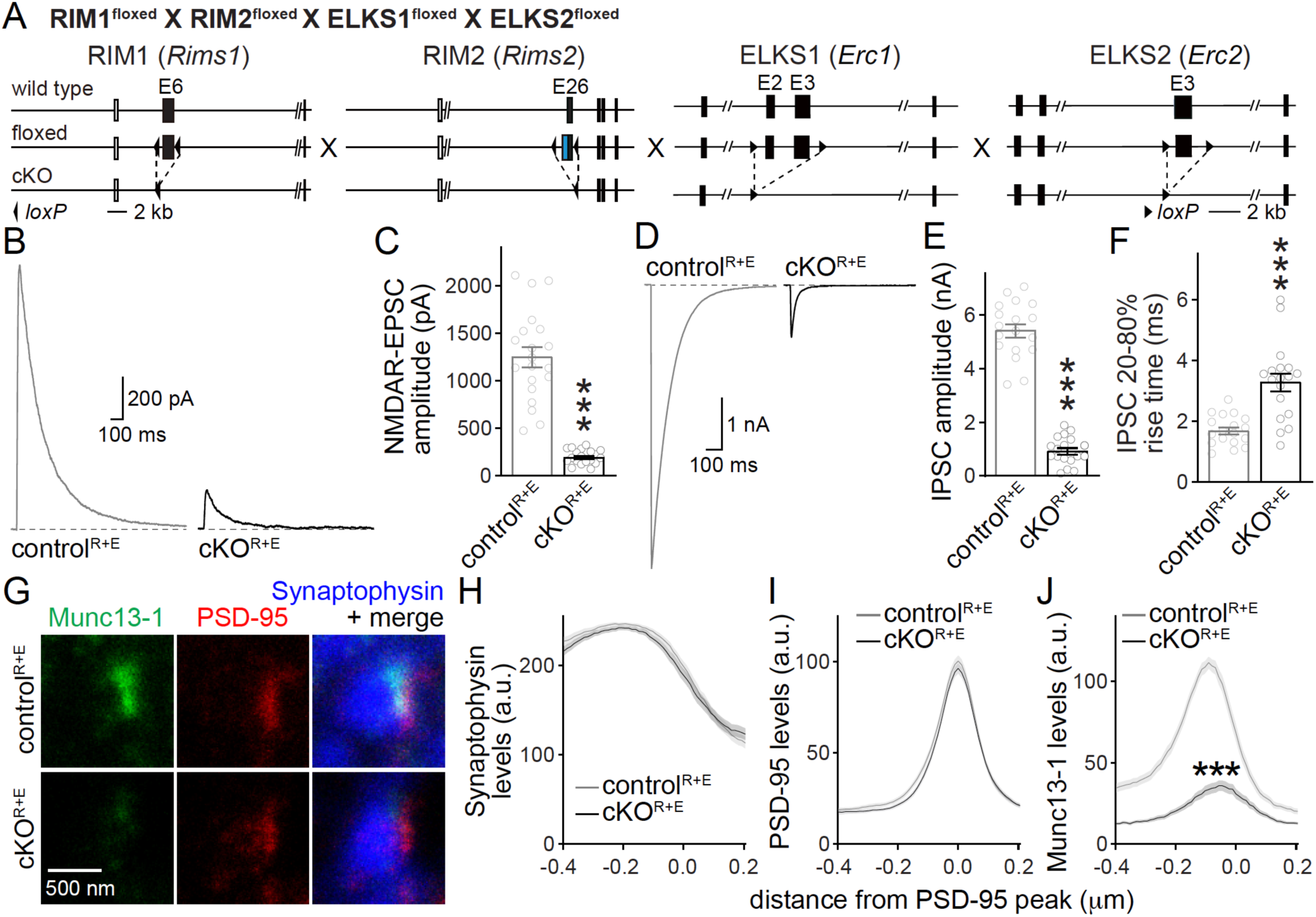
Action potential-evoked neurotransmitter release and Munc13 active zone levels after ablation of RIM+ELKS. (**A**) Targeting strategy for deletion of RIM1, RIM2, ELKS1 and ELKS2 in cultured hippocampal neurons. Neurons of mice with floxed alleles for all four genes were infected with Cre-expressing lentiviruses (to generate cKO^R+E^ neurons) or lentiviruses expressing a recombination-deficient version of Cre (to generate control^R+E^ neurons) as described (Tan et al., 2022; Wang et al., 2016). (**B, C**) Sample traces (B) and quantification (C) of EPSCs evoked by focal electrical stimulation, control^R+E^ 20 cells/3 cultures, cKO^R+E^ 19/3. (**D-F**) Sample traces (D) and quantification of amplitudes (E) and 20-80% rise times (F) of IPSCs evoked by focal electrical stimulation, 18/3 each. (**G-J**) Sample STED images (G) and quantification (H-J) of side-view synapses in cultured hippocampal neurons stained for Munc13-1 (imaged in STED), PSD-95 (imaged in STED), and Synaptophysin (imaged in confocal). Peak position and levels were analyzed in line profiles (600 nm x 200 nm) positioned perpendicular to the center of elongated PSD-95 structure and aligned to the PSD-95 peak, 60 synapses/3 cultures each. Data are mean ± SEM; \*\*\**P* < 0.001 as determined by Welch’s t-tests (C, E and F) or two-way ANOVA followed by Bonferroni’s multiple comparisons post-hoc tests (H-J). For assessment of Munc13-1 levels after Munc13 knockout using STED microscopy, see Fig. S1. For comparison of Munc13-1 expression by confocal microscopy or Western blotting in ELKS+RIM or Munc13 knockout neurons, see Figs. S2A-S2G.

We next evaluated Munc13-1 positioning and levels at the active zone using stimulated emission depletion (STED) superresolution microscopy (Figs. 1G-1J). We stained for Synaptophysin to mark the synaptic vesicle cloud (imaged by confocal microscopy), PSD-95 to mark the postsynaptic density (PSD, imaged by STED), and Munc13-1 (imaged by STED). We identified side-view synapses defined as a synaptic vesicle cluster with an elongated PSD-95 structure aligned at one side of the vesicle cloud as described before (Held et al., 2020; Nyitrai et al., 2020; Wong et al., 2018). We then quantified Munc13-1 in 200 × 600 nm line profiles that were positioned perpendicular to the PSD through the center of the PSD-95 signal. We found that Munc13-1 was removed efficiently from the active zone area of cKO^R+E^ synapses (Fig. 1J). For comparison, we pursued analyses of Munc13-1 levels at the active zone after ablation of Munc13. We used mice with floxed alleles for Munc13-1 and constitutive knockout alleles for Munc13-2 and Munc13-3 (Fig. S1A) (Augustin et al., 2001; Banerjee et al., 2022; Varoqueaux et al., 2002). In cultured neurons of these mice, Cre-expression removes Munc13-1 (cKO^M^) without the potential for compensation by Munc13-2 or −3 because these genes are knocked out constitutively, and control experiments were performed on the same cultures but with expressing a lentivirus with an inactive Cre (control^M^). We found that Munc13-1 was removed efficiently (Figs. S1B-S1D), with the leftover signal not distinguishable from background levels that are typically observed in this approach (Held et al., 2020; Nyitrai et al., 2020; Wong et al., 2018). When we compared Munc13-1 levels in cKO^R+E^ and cKO^M^ synapses, the remaining signal was somewhat higher in cKO^R+E^ synapses. These higher levels did not arise from a peak at the position of the active zone area (around −70 to −20 nm) (Tan et al., 2022; Wong et al., 2018). Instead, Munc13-1 levels appeared broadly increased, and the ratio of Munc13-1 at cKO^R+E^ vs. cKO^M^ synapses shifted upwards throughout the presynaptic bouton (Fig. S1F). In both types of neurons, Synaptophysin and PSD-95 signals remained unchanged (Figs. 1H, 1I, S1C, S1E), confirming that synapse assembly is not strongly affected because synaptic vesicle clusters are aligned with PSDs.

Because a small amount of remaining Munc13-1 could be detected in cKO^R+E^ neurons by STED microscopy (Figs. 1G-1J, S1F) and by Western blotting (Wang et al., 2016), we compared synaptic Munc13-1 levels in cKO^R+E^ and cKO^M^ neurons using confocal microscopy, in images taken from the same set of cultures that were analyzed by STED. We found that somewhat higher synaptic Munc13-1 signals, quantified as intensity within synaptophysin regions of interest (ROIs), remained at cKO^R+E^ synapses compared to cKO^M^ synapses (Figs. S2A-S2E). Similarly, a slight Munc13-1 band was detected in Western blotting of cKO^R+E^ neurons, but not in cKO^M^ neurons (Figs. S2F, S2G). We note that the remaining Munc13-1 signal in immunostainings detected in cKO^M^ neurons is likely mostly composed of antibody background in these experiments, as quantifications are done without background subtraction and noise levels of ∼25% are common (Wang et al., 2016). Altogether, our data indicate that some Munc13-1 might remain in nerve terminals of cKO^R+E^ synapses, but the remaining Munc13 is not efficiently concentrated in the active zone area apposed to the PSD, supporting previous work on RIM-mediated recruitment of Munc13 to the active zone (Andrews-Zwilling et al., 2006).

### Deletion of Munc13 in addition to RIM and ELKS further impairs synaptic vesicle release

To test whether Munc13-1 mediates the remaining release in cKO^R+E^ neurons, we crossed the conditional knockout mice for RIM1, RIM2, ELKS1 and ELKS2 to conditional Munc13-1 and constitutive Munc13-2 knockout mice (Fig. 2A). Cultured hippocampal neurons from these mice were infected with Cre-expressing or control lentiviruses at DIV5 to generate cKO^R+E+M^ and control^R+E+M^ neurons, respectively, to remove Munc13 in addition to RIM+ELKS (Figs. 2A, S2H). We used confocal and STED microscopy to analyze the localization and levels of Synaptophysin, PSD-95, RIM1 and Munc13-1 (Figs. 2B-2M). Both RIM1 and Munc13-1 were effectively removed from active zones of cKO^R+E+M^ neurons, while the distribution and levels of Synaptophysin and PSD-95 were unaffected (Figs. 2B-2I). To analyze synapse numbers, we used a custom-written code to perform automatic two-dimensional segmentation (Emperador-Melero et al., 2021; Held et al., 2020; Liu et al., 2018) for Synaptophysin object detection and quantified the density, intensity and area of Synaptophysin puncta. These parameters were indistinguishable between control^R+E+M^ and cKO^R+E+M^ neurons (Figs. 2J-2M).

**Figure 2.**
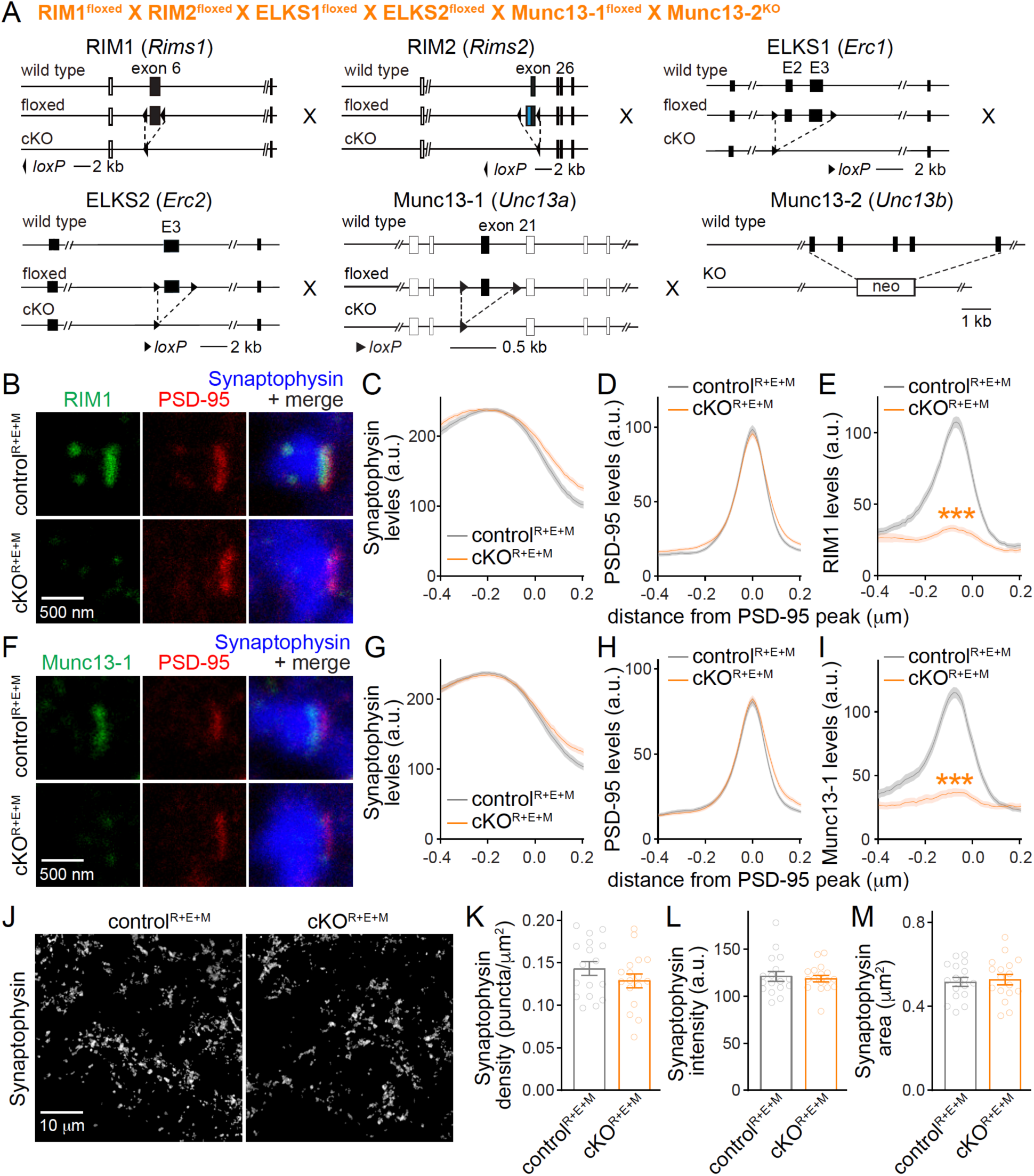
Simultaneous deletion of RIM, ELKS and Munc13 does not disrupt synapse formation. (**A**) Targeting strategy for simultaneous deletion of RIM1, RIM2, ELKS1, ELKS2, Munc13-1 and Munc13-2 in cultured hippocampal neurons (cKO^R+E+M^). Neurons were infected with Cre-expressing lentiviruses (to generate cKO^R+E+M^ neurons) or viruses expressing a recombination-deficient version of Cre (to generate control^R+E+M^ neurons). (**B-E**) Sample STED images (B) and quantification (C-E) of side-view synapses stained for RIM1 (STED), PSD-95 (STED), and Synaptophysin (confocal), control^R+E+M^ 65 synapses/3 cultures, cKO^R+E+M^ 66/3. (**F-I**) Same as B-E, but stained for Munc13-1, control^R+E+M^ 65/3, cKO^R+E+M^ 63/3. (**J-M**) Overview confocal images of anti-Synaptophysin staining (J) and quantification of Synaptophysin puncta density (K), intensity (L) and size (M); synaptophysin objects were detected using automatic two-dimensional segmentation, synaptophysin confocal images are from the experiment shown in B and C, 17 images/3 cultures each. Data are mean ± SEM; \*\*\**P* < 0.001 as determined by two-way ANOVA followed by Bonferroni’s multiple comparisons post-hoc tests (C-E, G-I), or unpaired two-tailed Student’s t-tests (K-M). For Munc13-1 levels tested by Western blotting, see Fig. S2H.

We next analyzed synaptic ultrastructure using high pressure freezing followed by transmission electron microscopy in the cultured neurons (Fig. 3), following established procedures (Tan et al., 2022; Wang et al., 2016). In these analyses, docked synaptic vesicles are defined as vesicles for which the electron density of the vesicular membrane merges with that of the presynaptic plasma membrane, and less electron-dense space cannot be detected between the two membranes. Simultaneous deletion of RIM, ELKS and Munc13 abolished vesicle docking (Figs. 3A, 3B) similar to RIM+ELKS ablation (Tan et al., 2022; Wang et al., 2016). While overall bouton size was unchanged (Fig. 3D), there were small decreases in vesicle numbers and mild increases for PSD-95 length in cKO^R+E+M^ neurons (Figs. 3C, 3E). This might be caused by homeostatic adaptations or by general roles of these proteins in synapse development, or be coincidental. Altogether, the morphological analyses establish that nerve terminals and synaptic appositions form and are overall ultrastructurally near-normal (apart from a loss of docked vesicles) despite the strong genetic manipulation with deletion of RIM1, RIM2, ELKS1, ELKS2, Munc13-1 and Munc13-2.

**Figure 3.**
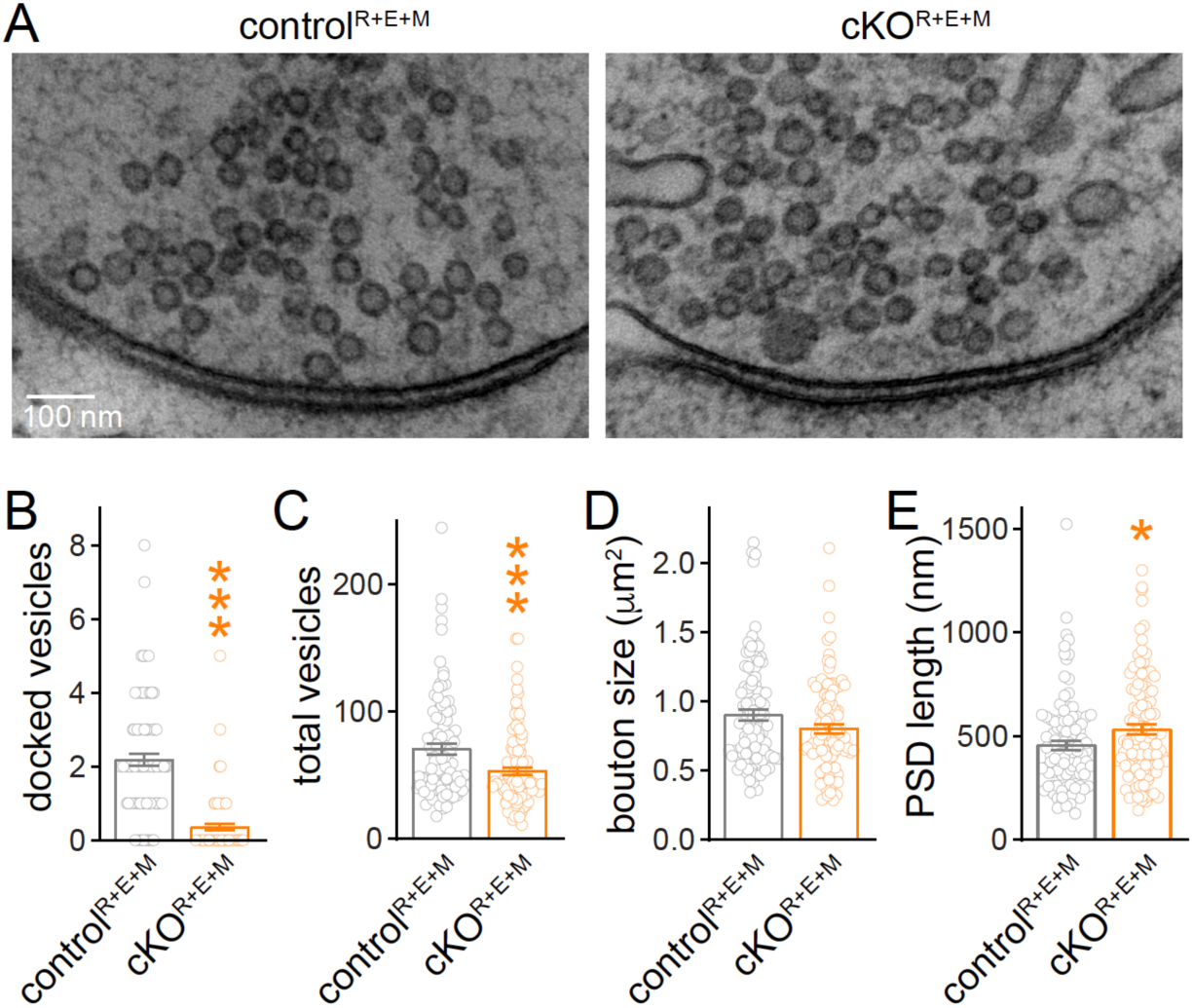
Synaptic ultrastructure after RIM+ELKS+Munc13 knockout. (**A-E**) Sample images (A) and analyses (B-E) of synaptic morphology of high-pressure frozen neurons analyzed by electron microscopy; docked vesicles (B), total vesicles (C), bouton size (D) and PSD length (E) per synapse profile are shown. Docked vesicles were defined as vesicles for which the electron densities of the vesicular and target membranes merge such that they are not separated by less electron dense space, control^R+E+M^ 99 synapses/2 cultures, cKO^R+E+M^ 100/2. Data are mean ± SEM; \**P* < 0.05, \*\*\**P* < 0.001 as determined by Welch’s t-tests (B and C), or by unpaired t two-tailed Student’s tests (D and E).

Using whole-cell recordings, we then assessed synaptic transmission in cKO^R+E+M^ neurons and corresponding controls. We first measured spontaneous vesicle release by assessing frequencies and amplitudes of miniature excitatory and inhibitory postsynaptic currents (mEPSCs and mIPSCs, respectively) in the presence of the sodium channel blocker tetrodotoxin. The frequencies of mEPSCs and mIPSCs were robustly decreased in cKO^R+E+M^ neurons, while their amplitudes remained unchanged (Figs. 4A-4F). Hence, vesicle release is impaired, but postsynaptic neurotransmitter detection is intact.

**Figure 4.**
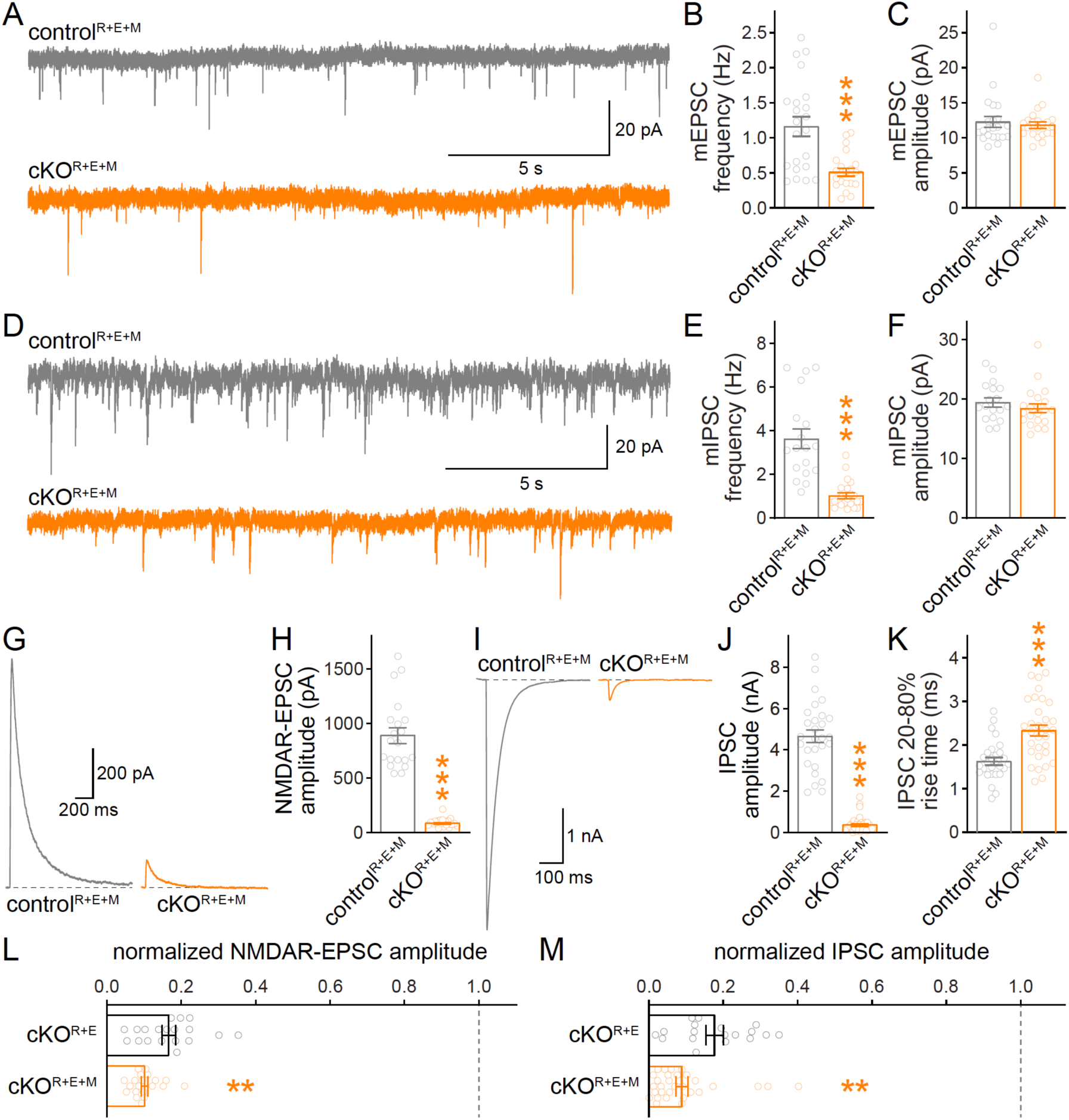
Neurotransmitter release is strongly impaired after RIM+ELKS+Munc13 ablation. (**A-C**) Sample traces (A) and quantification of mEPSC frequencies (B) and amplitudes (C), 22 cells/3 cultures each. (**D-F**) Sample traces (D) and quantification of mIPSC frequencies (E) and amplitudes (F), control^R+E+M^ 18/3, cKO^R+E+M^ 21/3. (**G, H**) Sample traces (G) and quantification (H) of EPSCs evoked by focal electrical stimulation, 20/3 each. (**I-K**) Sample traces (I) and quantification of amplitudes (J) and 20-80% rise times (K) of IPSCs evoked by focal electrical stimulation, control^R+E+M^ 28/3, cKO^R+E+M^ 31/3. (**L**) Comparison of EPSCs normalized to their own controls for cKO^R+E^ (absolute data from Fig. 1C) and cKO^R+E+M^ (from H) neurons, cKO^R+E^ 19/3, cKO^R+E+M^ 20/3. (**M**) Comparison of IPSCs normalized to their own controls for cKO^R+E^ (absolute data from Fig. 1E) and cKO^R+E+M^ (from J) neurons, cKO^R+E^ 18/3, cKO^R+E+M^ 31/3. Data are mean ± SEM; \*\**P* < 0.01, \*\*\**P* < 0.001 as determined by Welch’s t-tests (B, K and L), or Mann-Whitney tests (C, E, F, H, J and M).

We then measured single action potential-evoked release at cKO^R+E+M^ synapses (Figs. 4G-4K). The cKO^R+E+M^ neurons had very strong reductions in evoked release, both at excitatory (Figs. 4G, 4H) and inhibitory (Figs. 4I-4K) synapses compared to control^R+E+M^ neurons. To directly compare this reduction to that observed in cKO^R+E^ neurons (Figs. 1B-1E), we normalized the cKO PSC amplitudes to their own controls, which are genetically identical except for the expression of Cre recombinase. We then compared the normalized data across genotypes (Figs. 4L, 4M). This analysis established that for both excitatory and inhibitory synapses, knockout of Munc13 on top of RIM and ELKS led to a significantly stronger reduction in synaptic transmission. The small amount of remaining release might be due to the low amount of exon 21/22-deficient Munc13-1 that persists after conditional Munc13-1 knockout with this allele (Banerjee et al., 2022), to Munc13-1 that persists beyond 11 days of Cre expression, to Munc13-3 that was not deleted in the hextuple knockout mice, or to an alternative release pathway that does not depend on RIM, ELKS and Munc13. Altogether, however, the data establish that the remaining synaptic transmission after RIM and ELKS knockout depends on the presence of Munc13-1 and Munc13-2.

### Munc13 contributes to a remaining functional RRP after active zone disruption

Given the further reduction of synaptic transmission when Munc13 is ablated in cKO^R+E^ neurons, we analyzed vesicle priming and release probability in cKO^R+E+M^ neurons. The goal was to determine which release properties are controlled by Munc13 through comparison of these parameters with cKO^R+E^ neurons. We assessed the functional RRP at both excitatory and inhibitory synapses through the application of hypertonic sucrose, a method that has been broadly used to evaluate correlations between vesicle docking and priming (Imig et al., 2014; Rosenmund and Stevens, 1996; Wang et al., 2016; Zarebidaki et al., 2020). We detected robust reductions in the vesicle pool assessed by this method in cKO^R+E^ neurons (Figs. 5A-5D), but the functional RRP was not fully disrupted. Deletion of Munc13 on top of RIM and ELKS revealed an additional decrease, with an almost complete loss of releasable vesicles at excitatory cKO^R+E+M^ synapses and a >80% reduction at inhibitory cKO^R+E+M^ synapses (Figs. 5E-5H). Comparison of these two genotypes by normalizing each mutant to their corresponding control condition revealed that cKO^R+E+M^ neurons showed a significantly smaller RRP size (Figs. 5I, 5J) than cKO^R+E^ synapses in both synapse types. We conclude that the fusion-competence of vesicles that remains after active zone disruption by RIM+ELKS knockout is mediated by Munc13. Because Munc13 is not active zone-anchored and docked vesicles are barely detectable in cKO^R+E^ neurons, these vesicles are undocked, but likely associated with Munc13 away from the active zone.

**Figure 5.**
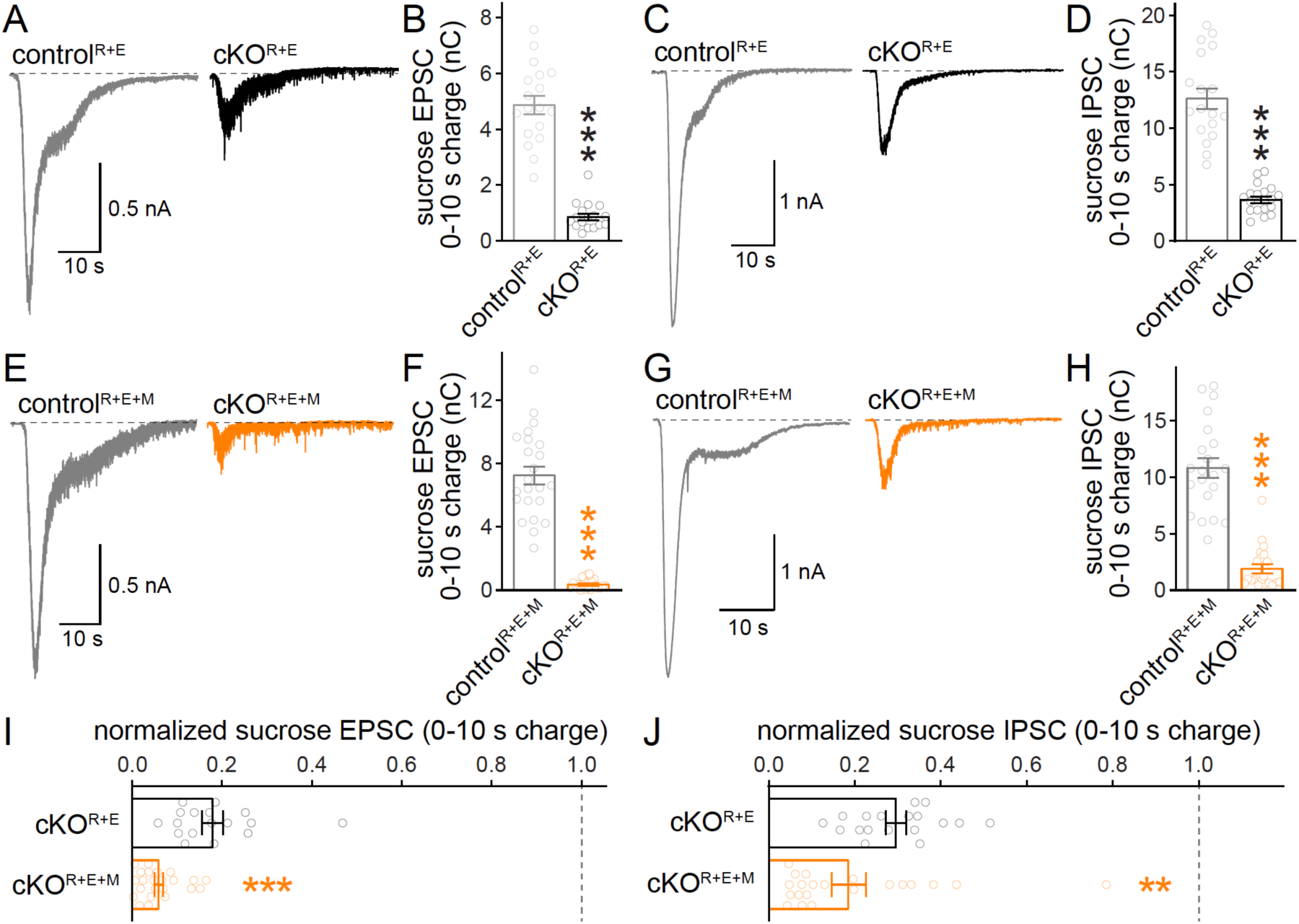
The remaining functional RRP in RIM+ELKS-deficient synapses depends on Munc13. (**A, B**) Sample traces (A) and quantification (B) of EPSCs triggered by hypertonic sucrose in control^R+E^ and cKO^R+E^ neurons, the first 10 s of the EPSC were quantified to estimate the RRP, control^R+E^ 18 cells/3 cultures, cKO^R+E^ 17/3. (**C, D**) As A and B, but for IPSCs, 18/3 each. (**E-H**) As for A-D, but for cKO^R+E+M^ neurons, F: 23/3 each, H: 21/3 each. (**I**) Comparison of EPSCs triggered by hypertonic sucrose normalized to their own controls for cKO^R+E^ (absolute data from B) and cKO^R+E+M^ (from F), cKO^R+E^ 17/3, cKO^R+E+M^ 23/3. (**J**) Comparison of IPSCs triggered by hypertonic sucrose normalized to their own controls for cKO^R+E^ (absolute data from D) and cKO^R+E+M^ (from H), cKO^R+E^ 18/3, cKO^R+E+M^ 21/3. Data are mean ± SEM; \*\**P* < 0.01, \*\*\**P* < 0.001 as determined by Mann-Whitney tests (B, F, H, I and J) or Welch’s t-test (D).

We finally used paired pulse ratios to monitor vesicular release probability p (Fig. 6). Paired pulse ratios are correlated inversely with p (Zucker and Regehr, 2002). In cKO^R+E^ neurons, p was strongly decreased, which was illustrated by robust increases in paired pulse ratios at short interstimulus intervals at both excitatory and inhibitory synapses (Figs. 6A-6D). In cKO^R+E+M^ neurons, paired pulse ratios were affected to an extent very similar to cKO^R+E^ neurons at both synapse types (Figs. 6E-6H). Direct comparison of genotypes through normalization further supported this conclusion as effects on p in cKO^R+E^ synapses and cKO^R+E+M^ synapses were indistinguishable (Figs. 6I, 6J). Hence, while the remaining Munc13 at cKO^R+E^ synapses is sufficient to maintain a small functional RRP, it does not enhance vesicular release probability of these vesicles.

**Figure 6.**
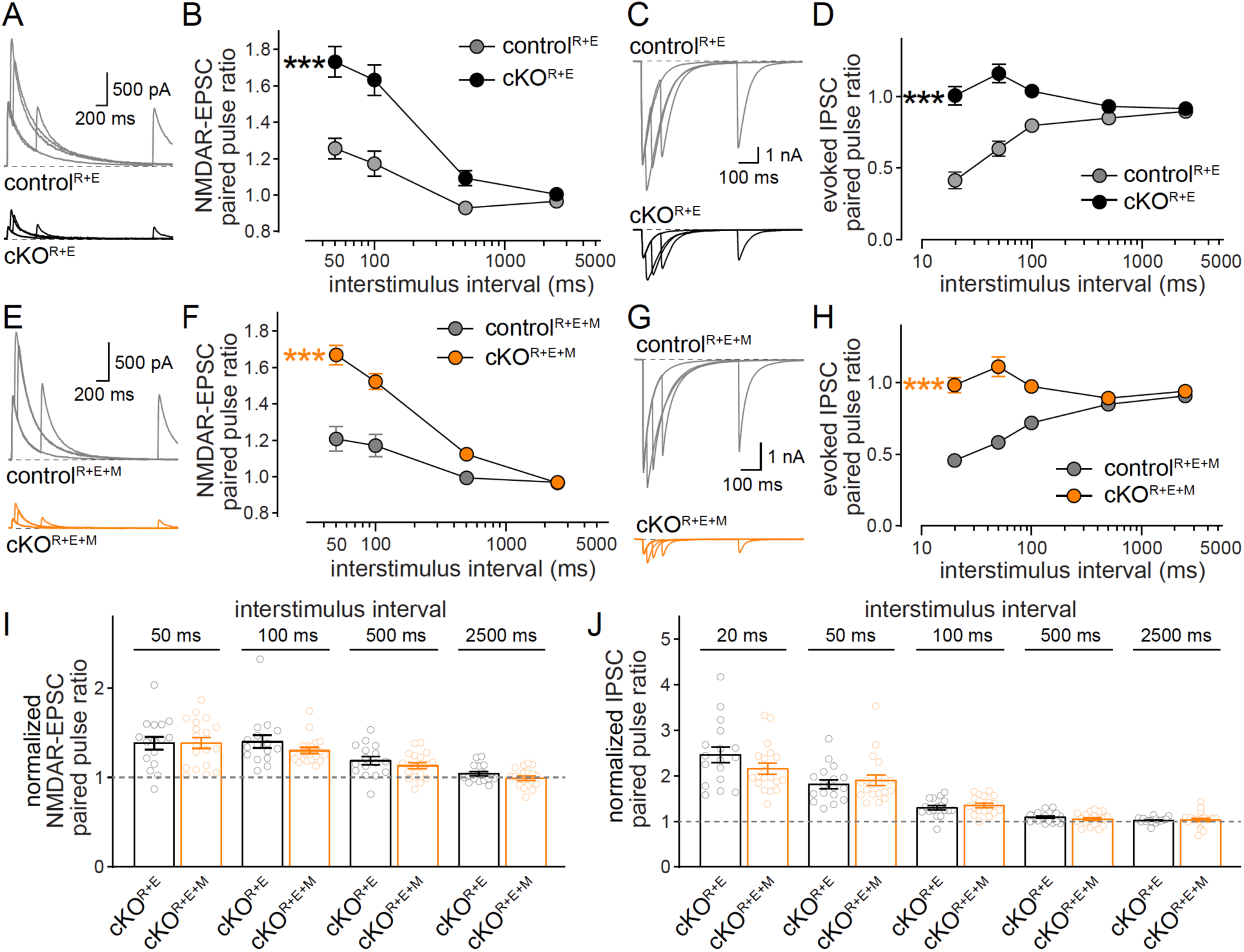
Vesicular release probability is not further affected by combined RIM+ELKS+Munc13 knockout. (**A, B**) Sample traces (A) and quantification (B) of EPSC paired pulse ratios in control^R+E^ and cKO^R+E^ neurons, control^R+E^ 15 cells/3 cultures, cKO^R+E^, 16/3. (**C, D**) As A and B, but for IPSCs (sample traces of 2500-ms intervals are not shown in C for simplicity), 17/3 each. (**E-H**) As for A-D, but for cKO^R+E+M^ neurons, F: 19/3 each, H: 19/3 each. (**I**) Comparison of EPSC paired pulse ratios across interstimulus intervals normalized to their own controls for cKO^R+E^ (absolute data from B) and cKO^R+E+M^ (from F), cKO^R+E^ 16/3, cKO^R+E+M^ 19/3. (**J**) Comparison of IPSC paired pulse ratios across interstimulus intervals normalized to their own controls for cKO^R+E^ (absolute data from D) and cKO^R+E+M^ (from H), cKO^R+E^ 17/3, cKO^R+E+M^ 19/3. Data are mean ± SEM; \*\*\**P* < 0.001 as determined by two-way ANOVA followed by Bonferroni’s multiple comparisons post-hoc tests (B, D, F and H), or by unpaired two-tailed Student’s t-tests (I and J).

## Discussion

We found previously that the functional RRP is not fully disrupted after ablating vesicle docking by simultaneous knockout of RIM and ELKS (Wang et al., 2016). Here, we show that fusion of these remaining RRP vesicles depends on Munc13. Even though Munc13-1 is not active zone-anchored after RIM and ELKS ablation, knocking out Munc13 in addition decreases the remaining releasable vesicles at excitatory and inhibitory hippocampal synapses. We conclude that Munc13 can render some vesicles fusogenic in the absence of RIM and ELKS. Our work adds the strongest compound knockout mutation to disrupt synaptic function with removal of six important active zone proteins to a growing body of literature using this approach. Beyond its relevance for mechanisms of neurotransmitter release, our work supports that the formation of synapses is remarkably resilient to even massive perturbations of presynaptic function and protein composition.

### Disrupting active zones to remove redundancy

Knockout studies on active zone gene families have defined multiple roles for these proteins in the neurotransmitter release process. While some functions and mechanisms, for example vesicle priming (Aravamudan et al., 1999; Augustin et al., 1999; Richmond et al., 1999; Varoqueaux et al., 2002), strongly depend on single proteins, other functions are redundant across protein families and hence more difficult to study mechanistically. This is particularly true for the scaffolding mechanisms that hold the active zone together and connect it with the target membrane and the vesicle cluster. These mechanisms are not well defined and models ranging from self-assembly of complexes with defined stoichiometries to phase separation based on multivalent low-affinity interactions have been proposed (Chen et al., 2020; Emperador-Melero and Kaeser, 2020; Südhof, 2012). Recent studies with compound mutants that ablate combinations of active zone protein families started to identify the required active zone scaffolds (Acuna et al., 2016; Kushibiki et al., 2019; Oh et al., 2021; Wang et al., 2016). We found previously that simultaneous deletion of RIM and ELKS in hippocampal neurons strongly disrupts active zone protein assemblies with loss of Munc13, RIM-BP, Piccolo, and Bassoon (Tan et al., 2022; Wang et al., 2016; Wong et al., 2018). This loss of active zone material causes a near-complete disruption of vesicle docking, and strongly impaired action potential-triggered release. Unexpectedly, however, some release persisted, either due to the presence of remaining release-competent but non-docked vesicles, or due to the rapid generation of release-competent vesicles in response to stimulation. The present study establishes that the transmitter release that remains after RIM and ELKS deletion relies on Munc13, as ablating Munc13 on top of RIM and ELKS robustly decreased the remaining functional RRP. We further found that synapse structure per se is resilient to this major genetic and functional perturbation, as synapses formed at normal densities and showed overall normal ultrastructure despite the removal of six important active zone genes via Cre recombination in the cultured neurons. Our work adds to a growing body of data demonstrating that neurotransmitter release, presynaptic Ca^2+^ entry, and active zone scaffolding are all dispensable for the formation of prominent types of central nervous system synapses (Held et al., 2020; Sando et al., 2017; Sigler et al., 2017; Verhage et al., 2000).

One way to interpret our data is to compare them with properties of fusion in other secretory pathways. Synaptic vesicle exocytosis after RIM and ELKS knockout has resemblance with secretion from chromaffin cells (Neher, 2018; Wang et al., 2016). In these cells, the release-ready pool of vesicles depends on Munc13, has relatively slower kinetics compared to synapses, but appears to not rely on the sequences of Munc13 that interact with RIM and on active zone scaffolds more generally (Betz et al., 2001; Man et al., 2015; Neher, 2018). The similarities of the remaining fusion after disrupting active zone assembly by knockout of RIM and ELKS are striking in that the release kinetics are slowed down (Wang et al., 2016), the remaining fusion depends on Munc13 (Figs. 3-6), and docking and the functional RRP are not fully correlated. Some of these similarities are also reminiscent of recent studies in C. elegans, where vesicle priming does not rely on the interaction of unc-13/Munc13 with unc-10/RIM, and possibly RIM itself (Liu et al., 2019). Altogether, these studies and our work indicate that disrupting the active zone through RIM+ELKS knockout renders synaptic vesicle exocytosis similar to chromaffin cell secretion. Notably, there are specific synapses with similar properties. For example, at hippocampal mossy fiber synapses, strong depolarizations lead to RRP estimates that are larger than the number of docked vesicles (Maus et al., 2020), supporting the model that RRP vesicles can be rapidly generated and released during such stimuli, perhaps similar to cKO^R+E^ synapses.

### Can Munc13 prime vesicles in the absence of RIM?

Our data reveal that the transmitter release that remains after active zone disruption upon RIM and ELKS deletion depends on Munc13. Thus, Munc13 can render vesicles fusion-competent in the absence of RIM and when Munc13 is not anchored at the active zone. This mechanism, however, is inefficient, as it only maintains a fraction of the functional RRP of a wild type synapse. An alternative model is that fusion-competence is rapidly generated and immediately followed by exocytosis during pool-depleting stimuli. We recently established that the functional RRP after RIM+ELKS knockout can be further boosted by re-expressing RIM zinc finger domains without enhancing vesicle docking (Tan et al., 2022). The zinc finger domain was co-localized with the vesicle cluster, was not concentrated at the active zone, and recruited Munc13 to the vesicle cluster. Hence, Munc13 can enhance vesicle fusogenicity through association with non-docked vesicles, at least when this association is generated artificially. Here, we show that the remaining fusion in RIM+ELKS knockouts depends on Munc13 (Figs. 4, 5), but that Munc13-1 is barely detectable at the active zone. However, some Munc13 is present at synapses. These data indicate that endogenous Munc13 can be near vesicles and enhance their fusogenicity even if Munc13 is not active zone-anchored. Hence, Munc13 on non-docked vesicles might mediate the remaining fusion through generation of a pool of vesicles that can be rapidly primed and released upon stimulation. Altogether, a model arises that the rate-limiting step for generation of a functional RRP is the presynaptic recruitment of Munc13, even if Munc13 is not fully anchored and activated at the active zone. Upon stimulation, roles of Munc13 in SNARE complex assembly and fusion can be rapidly executed.

Previous work established that RIM recruits Munc13 to active zones and activates it. This mechanism operates through binding of the RIM zinc finger to Munc13 C_2_A domains, which is necessary for rendering Munc13 monomeric and active in fusion (Andrews-Zwilling et al., 2006; Betz et al., 2001; Brockmann et al., 2020; Camacho et al., 2017; Deng et al., 2011; Dulubova et al., 2005; Lu et al., 2006). In the experiments presented here, the release remaining in the absence of RIM and ELKS depends on Munc13. This indicates that not all priming requires the RIM-mediated activation-mechanism of Munc13, consistent with previous observations that many but not all RRP vesicles are lost after RIM knockout (Han et al., 2011; Kaeser et al., 2011, 2012). An alternative mechanism could operate via bMunc13-2 (which lacks the C_2_A-domain that binds to RIM) and ELKS (Kawabe et al., 2017). While ELKS provides a Munc13-recruitment and priming mechanism that is independent of RIM, most ELKS is also removed in the RIM+ELKS knockout neurons (Kaeser et al., 2009; Liu et al., 2014; Wang et al., 2016), and only a very small subset of synapses relies on bMunc13-2 (Kawabe et al., 2017). Hence, this mechanism might be insufficient to explain the remaining release from RIM+ELKS knockout neurons. An alternative and perhaps more likely explanation is that not all C_2_A-domain containing Munc13 requires RIM. Monomeric, active Munc13 is in equilibrium with dimeric, inactive Munc13, and RIM shifts the equilibrium to the active form. Even in the absence of RIM, some Munc13 will be monomeric and available for assembling SNARE complexes, accounting for the vesicular exocytosis that remains in RIM+ELKS knockout neurons. Finally, this mechanism may not be restricted to docked vesicles, but vesicles associated with Munc13 may be amenable to release, explaining why some release persists in RIM+ELKS mutants despite the loss of active zone-anchored Munc13 and of a strong reduction in docked vesicles.

## Acknowledgements

We thank J. Wang for technical support, all members of the Kaeser laboratory for insightful discussions and feedback, and M. Verhage and J. Broeke for a MATLAB macro to analyze electron microscopic images. This work was supported by grants from the NIH (R01MH113349 and R01NS083898 to P.S.K.). We acknowledge the Neurobiology Imaging Facility (supported by a P30 Core Center Grant P30NS072030), and the Electron Microscopy Facility at Harvard Medical School.

## Author contributions

Conceptualization, C.T., N.B. and P.S.K.; Methodology, C.T., G.d.N., and C.Q.; Formal Analysis, C.T., G.d.N. and P.S.K.; Investigation, C.T., G.d.N. and C.Q.; Resources, C.T., C.Q., C.I. and N.B.; Writing-Original Draft, C.T. and P.S.K.; Writing-Review & Editing, C.T., C.Q., C.I., N.B. and P.S.K.; Supervision, P.S.K.; Funding Acquisition, P.S.K.

## Materials and Methods

### Resources table

| Reagent or resource | Designation | Source or reference | Identifiers | Additional information |
| --- | --- | --- | --- | --- |
| Genetic reagent - mouse | Rims1<sup>tm3Sud</sup>/J | (Kaeser et al., 2008) | RRID: IMSR_JAX:015832 |  |
| Genetic reagent - mouse | Rims2<sup>tm1.1Sud</sup>/J | (Kaeser et al., 2011) | RRID: IMSR_JAX:015833 |  |
| Genetic reagent - mouse | Erc1<sup>tm2.1Sud</sup>/J | (Liu et al., 2014) | RRID: IMSR_JAX:015830 |  |
| Genetic reagent - mouse | Erc2<sup>tm1.2Sud</sup>/J | (Kaeser et al., 2009) | RRID: IMSR_JAX:015831 |  |
| Genetic reagent - mouse | Munc13-1<sup>fl/fl</sup> | (Banerjee et al., 2022) | N/A |  |
| Genetic reagent - mouse | Munc13-2<sup>-/-</sup>; Unc13b<sup>tm1Rmnd</sup> | (Varoqueaux et al., 2002) | RRID_MGI: 2449706 |  |
| Genetic reagent - mouse | Munc13-3<sup>-/-</sup>; Unc13c<sup>tm1Bros</sup> | (Augustin et al., 2001) | RRID_MGI: 2449467 |  |
| Cell line | HEK293T cells | ATCC | RRID: CVCL_0063 |  |
| Recombinant DNA reagent | pFSW EGFP Cre | (Liu et al., 2014) | pHN131014, lab plasmid code (LPC): p009 |  |
| Recombinant DNA reagent | pFSW EGFP ΔCre | (Liu et al., 2014) | pHN131015, LPC: p010 |  |
| Antibody | rabbit anti-RIM | SySy | RRID: AB_887774, lab antibody code (LAC): A58 | Immunofluorescence (IF) (1:500) |
| Antibody | mouse anti-PSD-95 | NeuroMab | RRID: AB_10698024, LAC: A149 | IF (1:500) |
| Antibody | guinea pig anti-Synaptophysin | SySy | RRID: AB_1210382, LAC: A106 | IF (1:500) |
| Antibody | rabbit anti-Munc13-1 | SySy | RRID: AB_887733, LAC: A72 | IF (1:500); Western blot (WB) (1:1000) |
| Antibody | mouse anti-Synapsin | SySy | RRID: AB_2617071, LAC: A57 | WB (1:4000) |
| Antibody | goat anti-rabbit<br>Alexa Fluor 488 | Thermo Fisher | RRID: AB_2576217,<br>LAC: S5 | IF (1:200) |
| Antibody | goat anti-mouse<br>IgG2a Alexa Fluor<br>555 | Thermo Fisher | RRID: AB_1500824,<br>LAC: S20 | IF (1:200) |
| Antibody | goat anti-guinea<br>pig Alexa Fluor<br>633 | Thermo Fisher | RRID: AB_2535757,<br>LAC: S34 | IF (1:500) |
| Antibody | goat anti-mouse<br>peroxidase-<br>conjugated | MP Biologicals | RRID: AB_2334540;<br>LAC: S52 | WB (1:10000) |
| Antibody | goat anti-rabbit<br>peroxidase-<br>conjugated | MP Biologicals | RRID: AB_2334589;<br>LAC: S53 | WB (1:10000) |
| Software | MATLAB | MathWorks | RRID: SCR_001622;<br>https://www.mathworks.com/products/matlab.html |  |
| Software | Prism | GraphPad | RRID:<br>SCR_002798;<br>https://www.graphpad.com/scientific-software/prism |  |
| software | Fiji/ImageJ | NIH | RRID:<br>SCR_002285;<br>https://imagej.net/software/fiji/downloads |  |
| software | pClamp | Molecular<br>Devices | RRID:<br>SCR_011323;<br>https://www.moleculardevices.com/products/software/pclamp.html |  |
| algorithm | Custom MATLAB<br>code | (Liu et al., 2018,<br>2022) | doi.org/10.5281/zenodo.6388196 |  |

### Mouse lines

The quadruple homozygote floxed mice for RIM1αβ (RRID: IMSR_JAX:015832, (Kaeser et al., 2008)), RIM2αβγ (RRID: IMSR_JAX:015833, (Kaeser et al., 2011)), ELKS1α (RRID: IMSR_JAX:015830, (Liu et al., 2014)) and ELKS2α (RRID: IMSR_JAX:015831, (Kaeser et al., 2009)) were previously described (Wang et al., 2016). Exon 6 (E6) and 26 (E26) were flanked by loxP sites in the conditional RIM1αβ and RIM2αβγ knockout mice, respectively. Exons 2 (E2) and 3 (E3) were flanked by loxP sites in the conditional ELKS1α knockout mice, and exon 3 (E3) was flanked by loxP sites in conditional ELKS2α knockout mice. Floxed Munc13-1 mice (Banerjee et al., 2022) were crossed to constitutive knockout mice for Munc13-2 (Unc13b^tm1Rmnd^, RRID_MGI:2449706, (Varoqueaux et al., 2002)) and Munc13-3 (Unc13c^tm1Bros^, RRID_MGI:2449467, (Augustin et al., 2001)) to produce Munc13 homozygote knockout mice. Exon 21 (E21) was flanked by loxP sites in the conditional Munc13-1 knockout mice. Mice for simultaneous ablation of RIM1, RIM2, ELKS1, ELKS2, Munc13-1 and Munc13-2 were generated by crossing the corresponding conditional (RIM1αβ, RIM2αβγ, ELKS1α, ELKS2α, and Munc13-1) and constitutive (Munc13-2) knockout alleles to homozygosity.

#### Cell culture and lentiviral infection

Primary mouse hippocampal cultures were generated from newborn pups within 24 h after birth as described (Held et al., 2020; Tan et al., 2022; Wang et al., 2016) and cells from mice of both sexes were mixed. Mice were anesthetized on ice and the hippocampus was dissected out. Cells were dissociated and plated onto glass coverslips in tissue culture medium composed of Minimum Essential Medium (MEM) with 10% Fetal Select bovine serum (Atlas Biologicals FS-0500-AD), 2 mM L-glutamine, and 25 μg/mL insulin, 0.1 mg/mL transferrin, 0.5% glucose, 0.02% NaHCO_3_. Cultures were maintained in a 37 °C-tissue culture incubator, and after ∼24 h the plating medium was exchanged with growth medium composed of MEM with 5% Fetal Select bovine serum, 2% B-27 supplement (Thermo Fisher 17504044), 0.5 mM L-glutamine, 0.1 mg/mL transferrin, 0.5% glucose, 0.02% NaHCO_3_. At DIV3, depending on growth, 50% or 75% of the medium were exchanged with growth medium supplemented with 4 μM Cytosine β-D-arabinofuranoside (AraC) to inhibit glial cell growth. At DIV5, cultured neurons were infected with lentiviruses which were produced in HEK293T cells (RRID: CVCL_0063) by Ca^2+^ phosphate transfection. These lentiviruses expressed EGFP-tagged Cre recombinase (to generate cKO neurons) or a truncated, enzymatically inactive EGFP-tagged Cre protein (to generate control neurons). Expression in lentiviral constructs was driven by the human Synapsin promoter to restrict expression to neurons (Liu et al., 2014; Wang et al., 2016). Analyses were performed at DIV16-19.

#### Electrophysiology

Electrophysiological recordings in cultured hippocampal neurons were performed as described (Held et al., 2020; Tan et al., 2022; Wang et al., 2016) at DIV16-19. The extracellular solution contained (in mM): 140 NaCl, 5 KCl, 2 MgCl_2_, 1.5 CaCl_2_, 10 glucose, 10 HEPES-NaOH (pH 7.4, ∼300 mOsm). To avoid network activity induced by AMPA receptor activation, NMDAR-mediated excitatory postsynaptic currents (NMDAR-EPSCs) were measured to assess action potential-triggered excitatory transmission. For NMDAR-EPSCs, 6-Cyano-7-nitroquinoxaline-2,3-dione (CNQX, 20 μM) and picrotoxin (PTX, 50 μM) were present in the extracellular solution. Inhibitory postsynaptic currents (IPSCs) were recorded in the presence of D-amino-5-phosphonopentanoic acid (D-APV, 50 μM) and CNQX (20 μM) in the extracellular solution. Recordings were performed at room temperature (20 - 24 °C). Action potentials were elicited with a bipolar focal stimulation electrode fabricated from nichrome wire. Paired pulse ratios were calculated as the amplitude of the second PSC divided by the amplitude of the first at each interval from the average of 6 sweeps per cell and interval. The baseline value for the second PSC was taken immediately after the second stimulus artifact. For mEPSC and sucrose-induced EPSC recordings, TTX (1 μM), PTX (50 μM), and D-APV (50 μM) were added to the extracellular solution. For mIPSC and sucrose-induced IPSC recordings, TTX (1 μM), CNQX (20 μM), and D-APV (50 μM) were added to the extracellular solution. The RRP was estimated by application of 500 mM sucrose in extracellular solution applied via a microinjector syringe pump for 10 s at a rate of 10 μl/min through a tip with an inner diameter of 250 μm. mEPSCs and mIPSCs were identified with a template search followed by manual confirmation by an experimenter, and their frequencies and amplitudes were assessed during a 100-s recording time window after reaching a stable baseline (3-min after break-in). Glass pipettes were pulled at 2 - 5 MΩ and filled with intracellular solutions containing (in mM) for EPSC recordings: 120 Cs-methanesulfonate, 2 MgCl_2_, 10 EGTA, 4 Na_2_-ATP, 1 Na-GTP, 4 QX314-Cl, 10 HEPES-CsOH (pH 7.4, ∼300 mOsm) and for IPSC recordings: 40 CsCl, 90 K-gluconate, 1.8 NaCl, 1.7 MgCl_2_, 3.5 KCl, 0.05 EGTA, 2 Mg-ATP, 0.4 Na_2_-GTP, 10 phosphocreatine, 4 QX314-Cl, 10 HEPES-CsOH (pH 7.2, ∼300 mOsm). Cells were held at +40 mV for NMDAR-EPSC recordings and at −70 mV for evoked IPSC, mEPSC, mIPSC, sucrose EPSC and sucrose IPSC recordings. Access resistance was monitored and compensated to 3-5 MΩ, and cells were discarded if the uncompensated access exceeded 15 MΩ. Data were acquired at 5 kHz and lowpass filtered at 2 kHz with an Axon 700B Multiclamp amplifier and digitized with a Digidata 1440A digitizer. Data acquisition and analyses were done using pClamp10. For electrophysiological experiments, the experimenter was blind to the genotype throughout data acquisition and analysis.

#### STED and confocal imaging

Neurons were cultured on 0.17 mm thick 12 mm diameter coverslips. At DIV16-18, cultured neurons were washed two times with warm PBS, and fixed in 4% PFA in PBS for 10 min. After fixation, coverslips were rinsed twice in PBS, then permeabilized in PBS + 0.1% Triton X-100 + 3% BSA (TBP) for 1 h. Primary antibodies were diluted in TBP and stained for 24-48 h at 4 °C. The following primary antibodies were used: rabbit anti-RIM1 (1:500, RRID: AB_887774, A58), rabbit anti-Munc13-1 (1:500, RRID: AB_887733, A72), guinea pig anti-Synaptophysin (1:500, RRID: AB_1210382, A106), mouse anti-PSD-95 (1:500, RRID: AB_10698024, A149). After primary antibody staining, coverslips were rinsed twice and washed 3-4 times for 5 min in TBP. Alexa Fluor 488 (anti-rabbit, RRID: AB_2576217, S5), 555 (anti-mouse IgG2a, RRID: AB_1500824, S20), and 633 (anti-guinea pig, RRID: AB_2535757, S34) conjugated antibodies were used as secondary antibodies at 1:200 (Alexa Fluor 488 and 555) or 1:500 (Alexa Fluor 633) dilution in TBP, incubated for 24-48 h at 4 °C followed by rinsing two times and washing 3-4 times 5 min in TBP. Stained coverslips were post-fixed for 10 min with 4% PFA in PBS, rinsed two times in PBS + 50 mM glycine and once in deionized water, air-dried, and mounted on glass slides. STED images were acquired with a Leica SP8 Confocal/STED 3X microscope with an oil immersion 100x 1.44 numerical aperture objective and gated detectors as described (Emperador-Melero et al., 2021; Held et al., 2020; Tan et al., 2022; Wong et al., 2018). Images of 46.51 × 46.51 μm^2^ areas were scanned at a pixel density of 4096 × 4096 (11.358 nm/pixel). Alexa Fluor 633, Alexa Fluor 555, and Alexa Fluor 488 were excited with 633 nm, 555 nm and 488 nm using a white light laser at 2-5% of 1.5 mW laser power. The Alexa Fluor 633 channel was acquired first in confocal mode using 2x frame averaging. Subsequently, Alexa Fluor 555 and Alexa Fluor 488 channels were acquired in both confocal and STED modes. Alexa Fluor 555 and 488 channels in STED mode were depleted with 660 nm (50% of max power, 30% axial depletion) and 592 nm (80% of max power, 30% axial depletion) depletion lasers, respectively. Line accumulation (2-10x) and frame averaging (2x) were applied during STED scanning. Identical settings were applied to all samples within an experiment. Synapses within STED images were selected in side-view, defined as synapses that contained a synaptic vesicle cluster labeled with Synaptophysin and associated with an elongated PSD-95 structure along the edge of the vesicle cluster as described (Held et al., 2020; de Jong et al., 2018; Tan et al., 2022; Wong et al., 2018). For intensity profile analyses, side-view synapses were selected using only the PSD-95 signal and the vesicle signal for all experiments. A region of interest (ROI) was manually drawn around the PSD-95 signal and fit with an ellipse to determine the center position and orientation. A ∼1200 nm long, 200 nm wide rectangle was then positioned perpendicular to and across the center of the elongated PSD-95 structure. Intensity profiles from −400 nm (presynaptic) to +200 nm (postsynaptic) relative to the center of the PSD-95 signal were obtained for all three channels within this ROI. To align individual profiles, the PSD-95 signal only was smoothened using a moving average of 5 pixels, and the smoothened signal was used to define the peak position of PSD-95. All three channels (vesicle marker, test protein, and smoothened PSD-95) were then aligned to the PSD-95 peak position and averaged across images. For Figs. S2A-S2D, Munc13-1 levels were analyzed using Synaptophysin to define ROIs in the confocal images with Image J. For Figs. 2J-2M, ROI selection was performed using a previously described custom-written code to perform automatic two-dimensional segmentation (Emperador-Melero et al., 2021; Held et al., 2020; Liu et al., 2018, 2022) (code deposited at Zenodo: https://doi.org/10.5281/zenodo.6388196). After Synaptophysin object detection, the density, intensity and area of these objects were quantified. Analyses were performed on raw images without background subtraction, and adjustments were done identically across experimental conditions. Representative images were brightness and contrast adjusted to facilitate inspection, and these adjustments were made identically for images within an experiment. The experimenter was blind to the condition/genotype for image acquisition and analyses.

#### High-pressure freezing and electron microscopy

Neurons were cultured on 6 mm sapphire coverslips coated with matrigel. At DIV16-18, cultured neurons were frozen using a Leica EM ICE high-pressure freezer in extracellular solution containing (in mM): 140 NaCl, 5 KCl, 2 CaCl_2_, 2 MgCl_2_, 10 HEPES-NaOH (pH 7.4), 10 Glucose (∼300 mOsm) with PTX (50 μM), CNQX (20 μM) and D-AP5 (50 μM) added to block synaptic transmission. After freezing, samples were first freeze-substituted (AFS2, Leica) in anhydrous acetone containing 1% glutaraldehyde, 1% osmium tetroxide, 1% water. The process of freeze substitution was as follows: −90 °C for 5 h, 5 °C per h to −20 °C, −20 °C for 12 h, and 10 °C per h to 20 °C. Following freeze substitution, samples were Epon infiltrated, and baked for 48 h at 60 °C followed by 80 °C overnight before sectioning at 50 nm. For ultrathin sectioning, the sapphire coverslip was removed from the resin block by plunging the sample first in liquid nitrogen and followed by warm water several times until the sapphire was completely detached. The resin block containing the neurons was then divided into four pieces, and one piece each was mounted for sectioning. Ultrathin sectioning was performed on a Leica EM UC7 ultramicrotome, and the 50 nm sections were collected on a nickel slot grid (2 × 1 mm) with a carbon coated formvar support film. The samples were counterstained by incubating the grids with 2% lead acetate solution for 10 s, followed by rinsing with distilled water. Images were taken with a transmission electron microscope (JEOL 1200 EX at 80 kV accelerating voltage) and processed with ImageJ. The total number of vesicles, the number of docked vesicles per synapse profile, the area of the presynaptic bouton, and the length of the PSD were analyzed in each section using a custom-written Matlab code. Docked vesicles were defined as vesicles for which the electron densities of the vesicular membrane and the presynaptic plasma membrane merged such that the two membranes were not separated by less electron dense space. Bouton size was calculated from the measured perimeter of each synapse. Experiments and analyses were performed by an experimenter blind to the genotype.

#### Western blotting

At DIV15-19, cultured neurons were harvested in 20 μl 1x SDS buffer per coverslip and run on standard SDS-Page gels followed by transfer to nitrocellulose membranes. Membranes were blocked in filtered 10% nonfat milk/5% goat serum for 1 h at room temperature and incubated with primary antibodies (rabbit anti-Munc13-1, 1:1000, RRID: AB_887733, A72; mouse anti-Synapsin, 1:4000, RRID: AB_2617071, A57) in 5% nonfat milk/2.5% goat serum overnight at 4 °C, and HRP-conjugated secondary antibodies (1:10,000, anti-mouse, RRID: AB_2334540; anti-rabbit, RRID: AB_2334589) were used. For illustration in figures, images were adjusted for brightness and contrast to facilitate visual inspection, and the same adjustments were used for the entire scan.

#### Statistics

Statistics were performed in GraphPad Prism 9. Data are displayed as mean ± SEM, and significance is presented as * *P* < 0.05, ** *P* < 0.01, and *** *P* < 0.001. Parametric tests were used for normally distributed data (assessed by Shapiro-Wilk tests) or when sample size was n ≥ 30. Unpaired two-tailed Student’s t-tests were used for datasets with equal variance, or Welch’s unequal variances t-tests for datasets with unequal variance. For non-normally distributed data, Mann-Whitney tests were used. For paired pulse ratios, two-way ANOVA with Bonferroni’s post hoc tests was used. For STED side-view analyses, two-way ANOVA with Bonferroni’s post hoc tests was used on a 200 nm-window centered around the active zone peak. For all datasets, sample sizes and the specific tests used are stated in the figure legends.

**Figure S1.**
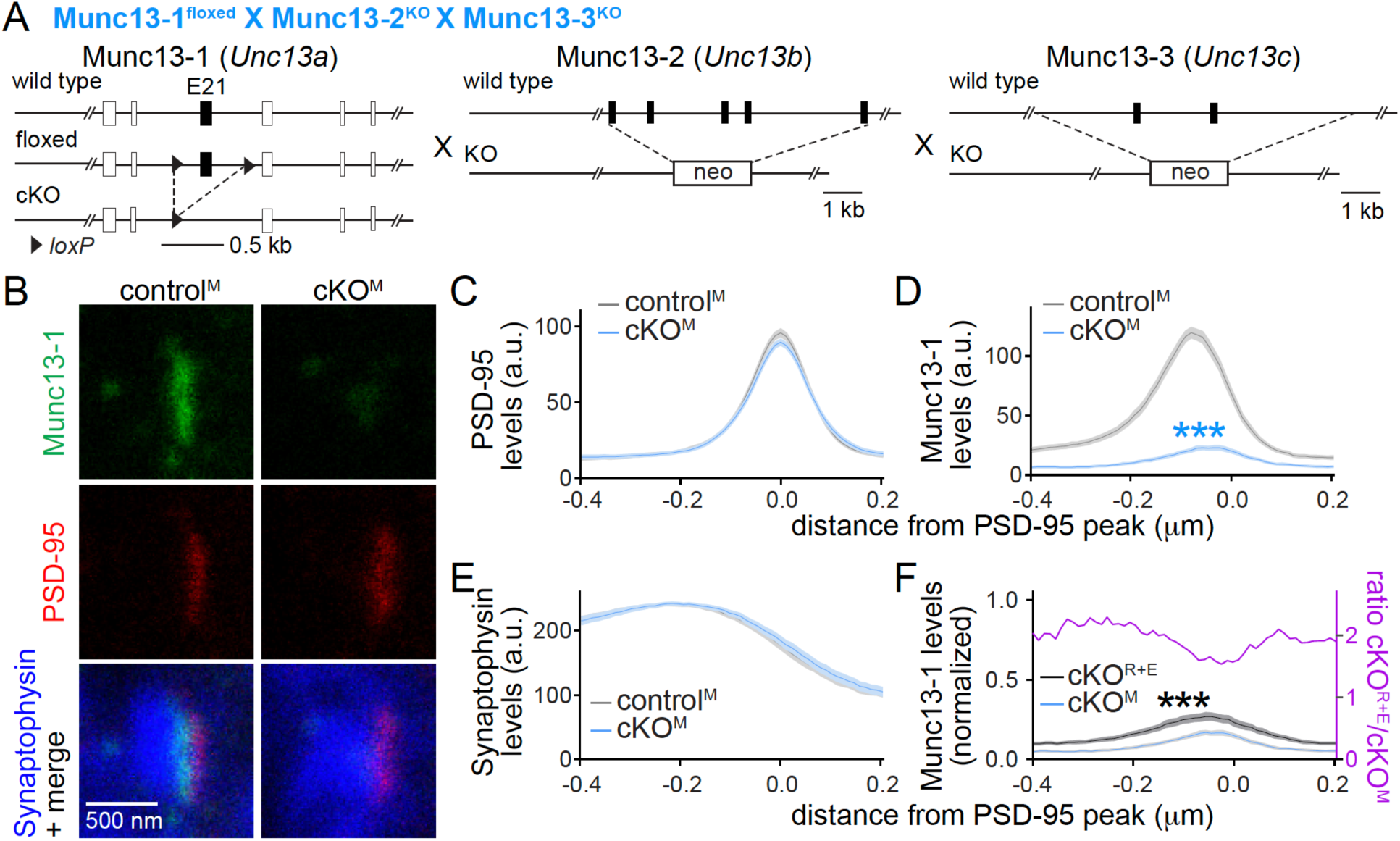
Assessment of Munc13-1 levels with STED microscopy. (**A**) Targeting strategy for deletion of Munc13-1, Munc13-2 and Munc13-3 in cultured hippocampal neurons (cKO^M^). Cultured hippocampal neurons of mice with floxed alleles for Munc13-1 (Banerjee et al., 2022) and constitutive knockout alleles for Munc13-2 (Varoqueaux et al., 2002) and Munc13-3 (Augustin et al., 2001) were infected with Cre-expressing lentiviruses (to generate cKO^M^ neurons) or lentiviruses expressing a recombination-deficient Cre (to generate control^M^ neurons). (**B-E**) Sample STED images (B) and quantification (C-E) of side-view synapses in cultured hippocampal neurons stained for Munc13-1 (STED), PSD-95 (STED), and Synaptophysin (confocal), 60 synapses/3 cultures each. (**F**) Comparison of Munc13-1 levels normalized to their own controls in cKO^R+E^ (absolute data from Fig. 1J) and cKO^M^ (from D) synapses, 60 synapses/3 cultures each. The purple line is the calculated ratio of normalized Munc13-1 levels from cKO^R+E^ to cKO^M^. Data are mean ± SEM; \*\*\**P* < 0.001 as determined by two-way ANOVA followed by Bonferroni’s multiple comparisons post-hoc tests (C-F).

**Figure S2.**
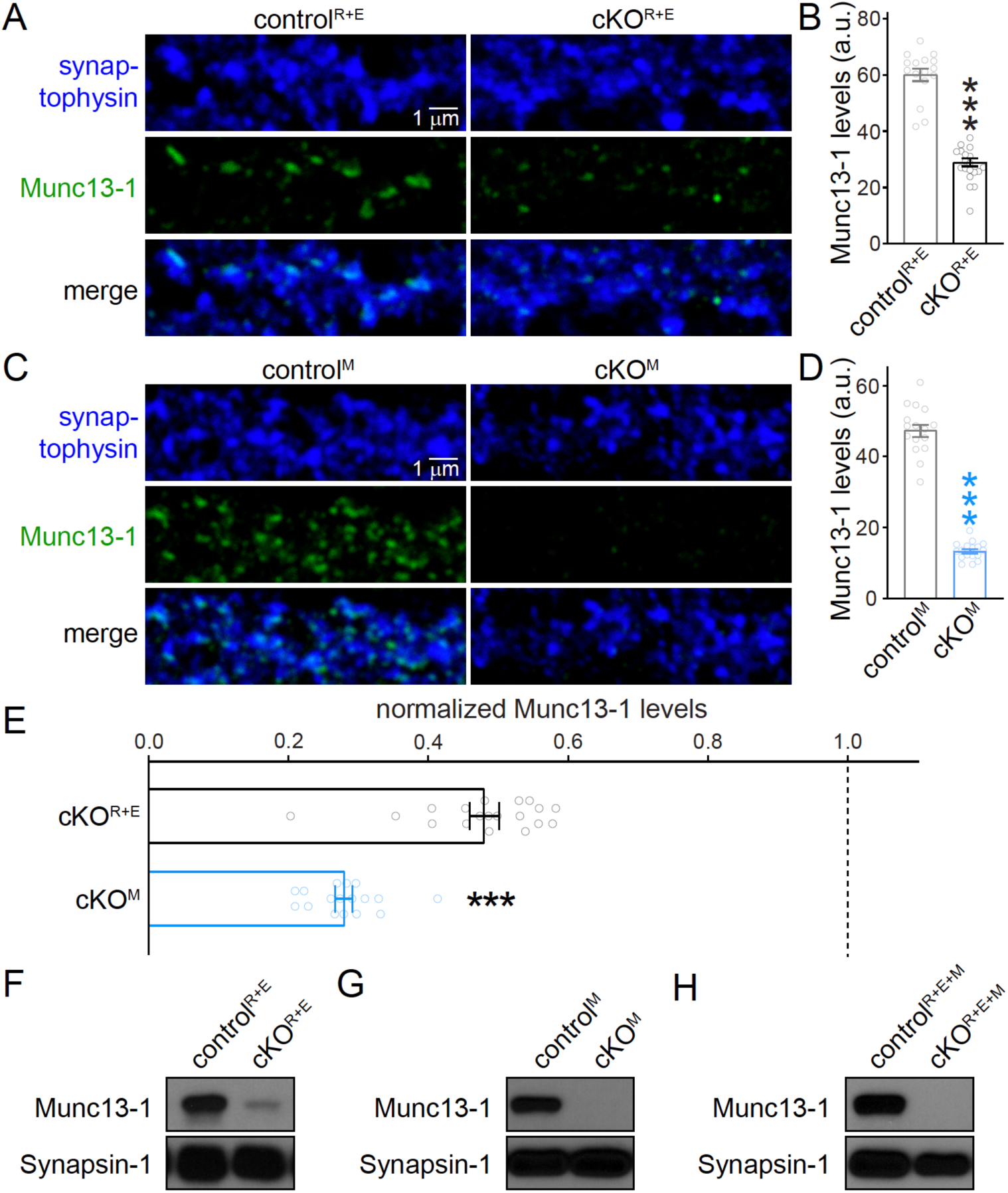
Assessment of Munc13-1 levels with confocal microscopy and Western blot. (**A, B**) Sample confocal images (A) and quantification (B) of Munc13-1 levels at control^R+E^ and cKO^R+E^ synapses. Synaptophysin staining was used to define ROIs, and levels of Munc13-1 were quantified within those ROIs, control^R+E^ 16 images/3 cultures, cKO^R+E^, 19/3. (**C, D**) As in A and B, but for cKO^M^ neurons, control^M^ 16/3, cKO^M^, 17/3. (**E**) Comparison of Munc13-1 levels normalized to their own controls for cKO^R+E^ (absolute data from B) and cKO^M^ (from D), cKO^R+E^ 19/3, cKO^M^ 17/3. (**F-H**) Assessment of Munc13-1 expression by Western blot of homogenates of cultured control^R+E^ and cKO^R+E^ neurons (F), control^M^ and cKO^M^ neurons (G) or control^R+E+M^ and cKO^R+E+M^ neurons (H) using anti-Munc13-1 antibodies and anti-Synapsin antibodies as loading controls. The results shown here are from ≤ 1 min exposure to film. The very low levels of an alternative variant of Munc13-1 left in Munc13-1 cKO neurons, described in Fig. S2 of (Banerjee et al., 2022), is not detected in these short exposures. Data are mean ± SEM; \*\*\**P* < 0.001 as determined by Welch’s t-test (D), or Mann-Whitney tests (B and E).

